# Anemia and tissue hypoxia are major determinants of malarial hypelactatemia

**DOI:** 10.1101/2024.10.22.619641

**Authors:** Athina Georgiadou, Chae Eun Yoon, Huanghehui Yu, William H. Pearson, Aika Ueno, Stefan Ebmeier, the SCRIPT Study Investigators, Gerald J. Larrouy-Maumus, Aubrey J. Cunnington

## Abstract

Hyperlactatemia, a key marker of severe malaria, is closely linked to increased mortality, though the exact mechanisms remain unclear. It may result from increased lactate production due to tissue hypoxia or reduced lactate clearance from organ dysfunction. This study used *Plasmodium yoelii* 17XL (Py17XL) murine model of severe malaria, which closely mimics hyperlactatemia seen in human cases, to investigate the contributions of severe anemia and infection-related organ dysfunction to hyperlactatemia. Non-infectious anemia models were also included for comparison. Anemia was found to elevate lactate in both malaria-infected and non-infectious models, but Py17XL infected mice showed higher lactate levels, indicating that anemia alone doesn’t fully explain hyperlactatemia. Evidence of tissue hypoxia, particularly in the liver, kidney, and gut, was seen with hypoxyprobe staining and upregulated hypoxia-inducible factor 1-alpha (HIF-1α), suggesting that hypoxia drives increased glycolysis and lactate production. Impaired lactate clearance may also play a role, as infected mice showed signs of liver and kidney dysfunction accompanied by reduced clearance of ^13^C_3_-labeled sodium-L-Lactate. Whole blood transfusion combined with artesunate significantly improved lactate clearance compared to artesunate alone, underscoring the importance of addressing anemia in treatment. A link between intestinal damage and hyperlactatemia was suggested by correlations between trefoil factor 3 (TFF3), a marker of gut injury, and lactate levels in human samples. Our findings highlight the multifactorial origin of hyperlactatemia in malaria, driven primarily by anemia and tissue hypoxia, pointing to the need for therapies targeting both aspects to reduce mortality in severe cases.

## Introduction

Malaria remains one of the world’s most prevalent infectious diseases, with 249 million new cases and 608,000 deaths reported in 2022, making it the leading vector-borne disease globally (1). Despite advances in eradication efforts, challenges persist due to the limited efficacy of available vaccines and the emergence of *Plasmodium* resistance to all current antimalarial drugs (2). Artemisinin-based combination therapy (ACT) remains the gold standard for treating malaria, however, some patients still succumb to the disease, and research has yet to identify life-saving adjunctive therapies (3). Severe malaria presents a variety of complications, including cerebral malaria, severe malarial anemia, acute respiratory distress syndrome, acute kidney injury, and metabolic disturbances like hyperlactatemia. Hyperlactatemia, characterized by blood lactate levels exceeding 5 mM, is a critical marker of severe malaria and is strongly associated with increased mortality (4, 5). Despite its close association with mortality the reasons for excess lactic acid accumulation and the mechanisms linking this to death are not well defined.

Lactic acidosis in malaria can be driven by an increase in lactate production, a reduction in lactate clearance or a combination of both (6). The rise in lactate production can stem from various sources. Parasite sequestration and reduced deformability of uninfected red blood cells may block blood vessels and reduce microcirculation, leading to localized tissue hypoxia, thus promoting anaerobic glycolysis (7). Furthermore, the destruction of both infected and uninfected red blood cells (RBCs) in malaria results in anemia, reducing the blood’s oxygen-carrying capacity and causing systemic hypoxia (8). Malaria induces a strong immune response, during which activated immune cells shift to aerobic glycolysis to meet their increased demand for ATP and essential metabolic intermediates needed for rapid proliferation. In these conditions, glycolysis is upregulated, resulting in an accelerated breakdown of glucose into pyruvate which is subsequently converted to lactate by lactate dehydrogenase A (LDHA) (9-11). Intraerythrocytic parasites, which depend on host’s glucose for energy, can produce lactate as a by-product (12) while a recent study suggested that when infected by *P. falciparum*, the RBC’s glycolytic rate can surge up to 100 times its normal level which may also contribute to the development of lactic acidosis and hypoglycemia in severe malaria (13). The liver and kidneys are responsible for 70% and 30% of lactate clearance respectively through gluconeogenesis (14). Liver dysfunction is a feature of both severe and uncomplicated malaria usually presenting as tissue damage, hemozoin deposits, white blood cell infiltration and elevated liver enzymes in blood [such as aspartate aminotransferase (AST) and alanine aminotransferase (ALT)](15). Acute kidney injury (AKI) is characterized by a sudden and significant decline in renal function and is one of the most critical complications of severe *P. falciparum* infection, which has consistently been linked to increased mortality rates in affected patients (16). Acute kidney injury (AKI) in malaria can result in the death of gluconeogenic cells within the renal cortex and a decrease in urine output, both of which hinder the body’s ability to clear lactate (6). While both liver and kidney have been suggested to exhibit reduced lactate clearance in severe malaria due to dysfunction (17), direct evidence supporting this remains limited.

Tissue hypoxia, often thought to be a major contributor to malaria-associated hyperlactatemia, arises when oxygen delivery is inadequate to meet metabolic demands (18). In severe malaria, hyperlactatemia is linked to poor tissue perfusion, caused by parasite sequestration in the microvasculature, anemia, and/or hypovolemia (19, 20). However, the underlying causes of lactic acidosis may vary based on malaria phenotypes (21). In cerebral malaria, a syndrome associated with extensive parasite sequestration, hyperlactatemia may result primarily from microvascular obstruction by parasites, impairing oxygen delivery. In contrast, in severe malarial anemia, it may stem from reduced oxygen-carrying capacity, forcing tissues into anaerobic metabolism and lactate overproduction (21). Although, the exact mechanisms by which anemia contributes to hyperlactatemia are not fully understood (22). Here we aimed to determine the contribution of severe malarial anemia to hyperlactatemia and reveal the mechanisms underlying this association.

We have previously discovered that mice infected with *P*.*yoelii* 17XL not only develop high blood lactate levels similar to those seen human severe malaria but also have the greatest blood transcriptome similarity to the human hyperlactatemia phenotype without the feature of sequestration (23). Here we use this severe malaria model together with a model of self-resolving malaria infection (*P*.*yoelii* 17XNL) and two models of non-infectious anemia (intravascular haemolysis induced by phenylhydrazine treatment (24) and extravascular haemolysis induced by anti-TER119 treatment (25)) to evaluate the relative contributions of anemia and tissue hypoxia to lactate production, and infection-induced impairment of lactate clearance, as possible mechanisms contributing to hyperlactatemia.

## Results

### Anemia is one of the main contributors to hyperlactatemia

To investigate the role of anemia in hyperlactatemia, four mouse models were employed: two malaria models and two anemia models (Fig.1A). In Py17XL infected mice, peak parasitemia, lactate levels, and nadir hematocrit occurred between days 5–7 post-infection, whereas Py17XNL infected mice reached peak parasitemia and lowest hematocrit around days 14–16 post-infection, after which they were expected to start clearing parasites and recovering. PHZ treated mice developed peak anemia three days after the first PHZ dose, and anti-TER119-treated mice reached peak anemia four days post the first anti-TER119 dose.

**Figure 1.**
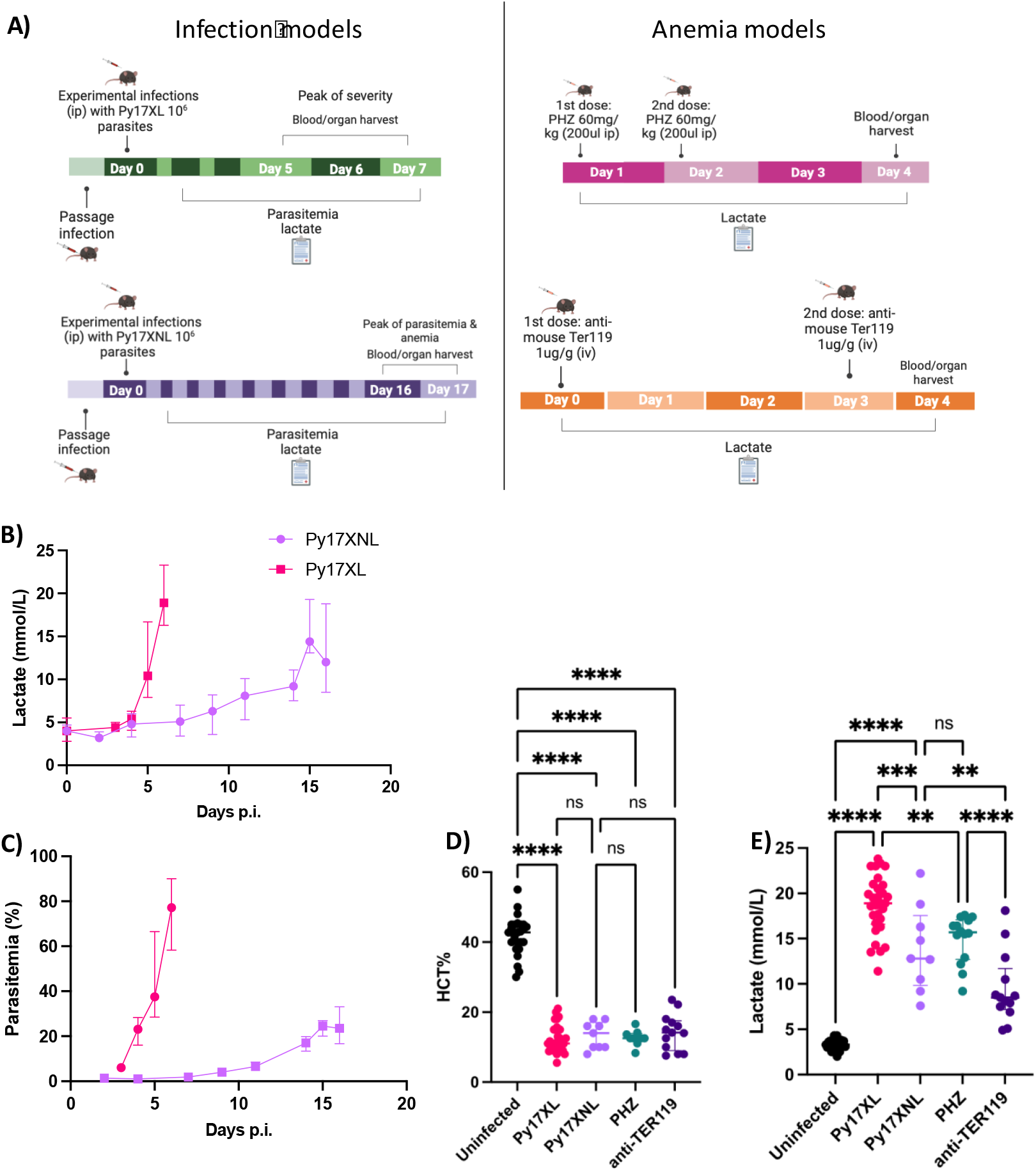
Anemia is one of the main contributors to high lactate. A) Malaria infection (Py17XL and Py17XNL) and anemia (PHZ and anti-TER119) mouse models timeline representation (figure created using Biorender). B) Lactate (mmol/L) levels comparison between Py17XL (n=9) and Py17XNL (n=9). C) Parasitemia (%) comparison between Py17XL (n=9) and Py17XNL (n=9). D) Hematocrit (%) levels at day of culling, E) Lactate (mmol/L) levels at day of culling. Uninfected n=25, Py17xl n=30, Py17XNL=9, PHZ n=14, anti-TER119 n=13. Error bars show median with interquartile range. One-way ANOVA *p-value* <0.0001, *p-values* for Tukey’s multiple comparison test: <0.0001= ****, 0.0005=***, <0.001=**. Eight to ten-week-old wild-type female C57BL/6J mice were used in all experiments. Py17XL: *P. yoelii* 17XL infected mice, Py17XNL: *P. yoelii* 17XNL infected mice, PHZ: phenylhydrazine treated mice, anti-TER119: anti-TER119 treated mice

When comparing the two malaria mouse models, we observed that both developed hyperlactatemia as parasitemia rose and hematocrit decreased, but the magnitude and progression varied. Py17XL infected mice had a median lactate level of 19 mmol/L (IQR = 17–22 mmol/L) and a median parasitemia of 77% (IQR = 60–89%) at six days post-infection, compared to Py17XNL-infected mice, which had a median lactate of 12 mmol/L (IQR = 8.9–18 mmol/L) and a median parasitemia of 24% (IQR = 17–29%) at 16 days post-infection (Fig.1B, C). Hematocrit levels showed no significant differences across the infection and anemia models, though all were significantly lower than in control, uninfected mice (Fig.1D).

Evaluating lactate levels on the day of hematocrit measurement, we found that both malaria infected and anemia models exhibited markedly elevated lactate, but Py17XL infected mice showed significantly higher lactate levels than the other conditions (Fig.2E). Since both the malaria and non-infectious anemia models all developed similar levels of anemia and high lactate, these results indicate that anemia is the major driver of hyperlactatemia, but additional mechanisms must contribute to the particularly high lactate seen in Py17XL infection.

**Figure 2.**
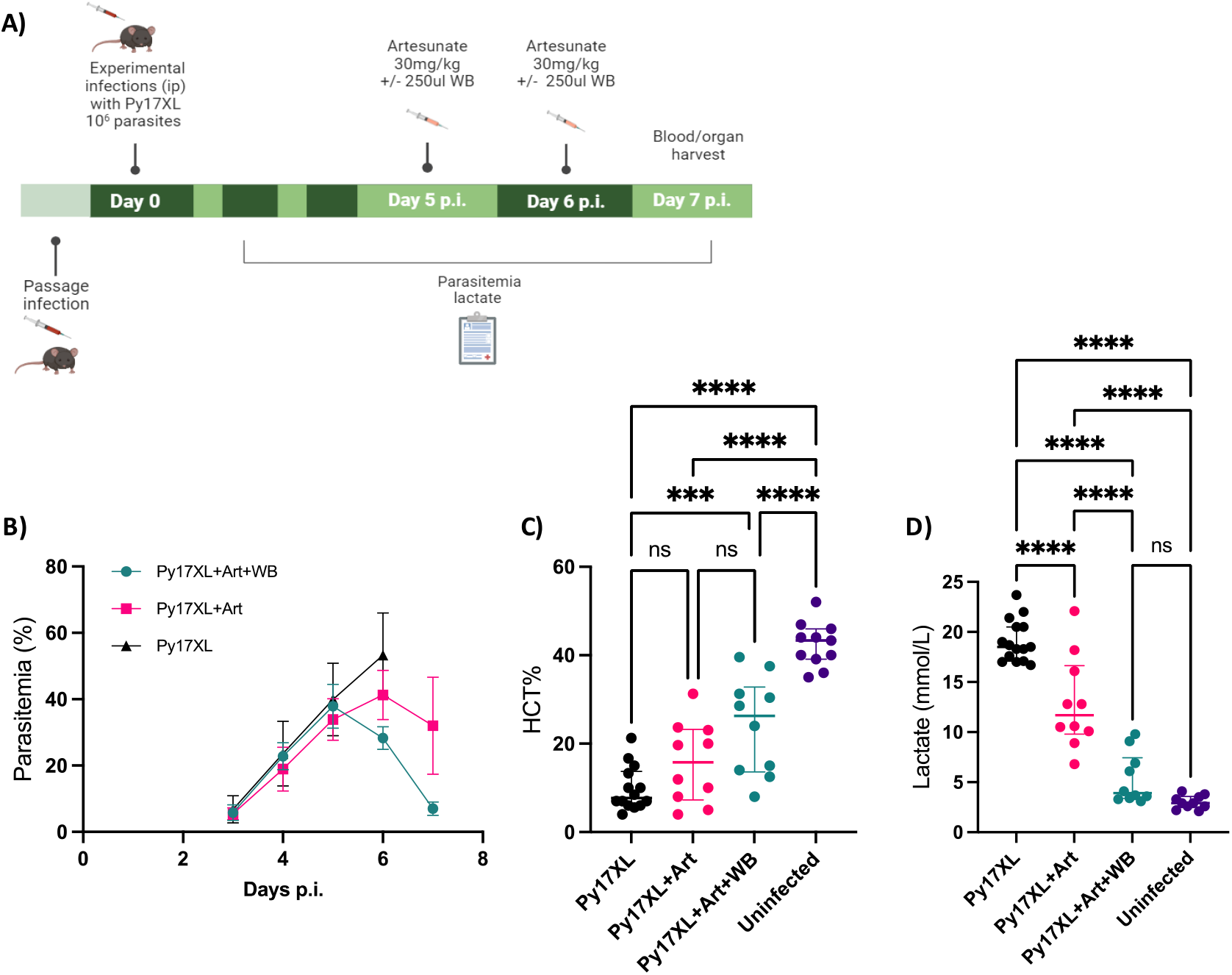
Combined treatment of Artesunate and whole blood transfusion improves resolution of lactate relative to Artesunate alone in Py17XL infected mice. A) Timeline of Py17XL infection and treatment with artesunate or artesunate plus whole blood starting at day 5 post infection (figure created using Biorender). B) Parasitemia (%) course over the days post infection, Py17XL n=12, Py17XL+Art n=10, Py17XL +Art + WB n=10, Error bars show mean with SD, C) Hematocrit (%) at the day of culling. One-way ANOVA *p-value* <0.0001, p-values for Tukey’s multiple comparison test: <0.0001= ****, 0.0003=***. D) Lactate (mmol/L) at the day of culling. One-way ANOVA *p-value* <0.0001, p-values for Tukey’s multiple comparison test: <0.0001= ****, Py17XL n=12, Py17XL+Art n=10, Py17XL +Art + WB n=10, Uninfected n=10. Error bars show median with interquartile range. Eight to ten-week-old wild-type female C57BL/6J mice were used in all experiments. Py17XL: *P. yoelii* 17XL infected mice, Art: artesunate, WB: whole blood.

### The combined treatment of artesunate and whole blood transfusion accelerates lactate clearance compared to artesunate alone in Py17XL infected mice

Having identified anemia as a major contributor to hyperlactatemia, we next evaluated the impact of combined whole blood transfusion and artesunate treatment versus artesunate alone on Py17XL infected mice, mimicking the approaches to treatment that might be taken in patients with severe malaria. Treatment was administered on days 5 and 6 post-infection, when the mice had already developed high parasitemia and elevated lactate levels (Fig.2A). At day 5 post-infection, before treatment, there were no significant differences in parasitemia among three Py17XL-infected groups: those treated with artesunate plus whole blood (combined treatment), those receiving artesunate alone, and untreated mice (Fig.2B). As expected, artesunate treatment began to reduce parasitemia within 24 hrs of initiation (Fig 2B). All infected groups had significantly lower hematocrit compared to uninfected mice (Fig.2C). Hematocrit levels, measured either on day 7 or at the humane endpoint (for untreated Py17XL), were significantly increased in the combined treatment group (*p-value* = 0.0003), but not the artesunate only group, compared to untreated mice (Fig.2C). Lactate levels, measured either on day 7 or at the humane endpoint (for untreated Py17XL), were significantly higher in untreated mice compared to both treatment groups suggesting that artesunate treatment can lead to the reduction of lactate (Fig.2D). However, importantly, lactate levels were significantly lower (and in many cases returned to normal range) in the mice receiving combined artesunate and whole blood transfusion compared to those treated with artesunate alone (Fig.2D). These findings suggest that the combination of artesunate and whole blood transfusion results in a faster resolution of anemia and lactate compared to artesunate alone, confirming that anemia causes hyperlactatemia.

### Tissue hypoxia in Py17XL infected mice

To investigate the localisation of tissue hypoxia in Py17XL infected mice compared to uninfected mice we administered Hypoxyprobe (Hypoxyprobe Inc, USA) *in vivo* on the day that infected mice reached humane endpoint in comparison to uninfected control mice. Among the different organs examined (liver, kidney, brain, spleen, gut, tail snip, heart and skeletal muscles) the liver, kidney, and gut of Py17XL infected mice showed prominent Hypoxyprobe staining (Fig.3).

**Figure 3.**
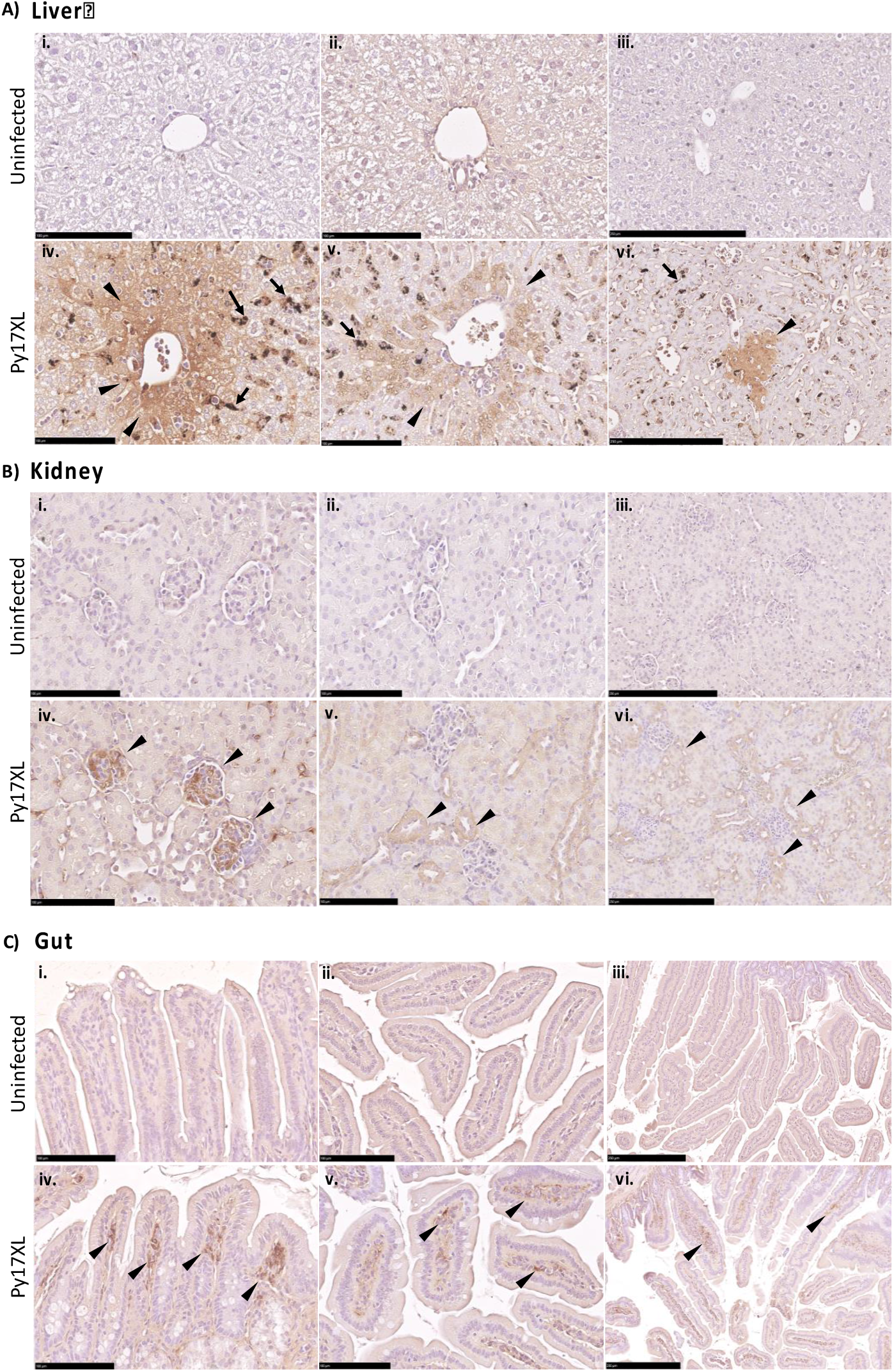
Localised hypoxia in Py17XL infected mice. Hypoxyprobe staining in A) liver, B) kidney and C) gut (small intestine) tissues. Arrowheads point towards hypoxyprobe staining. Arrows in the liver of Py17XL infected mice point towards hemozoin deposits. Images representative of control (n = 4) and Py17XL infected mice (n = 4). Scale bar for i, ii, iv and v: 100 μm, scale bar for iii and vi: 250 μm. Eight to ten-week-old wild-type female C57BL/6J mice were used in all experiments.

In the liver, no hypoxyprobe staining was detected in any of the uninfected samples (Fig.3A.i, ii, iii) whilst numerous patches of staining were seen in liver tissue of Py17XL infected mice (Fig.3A.iv, v, vi). Hypoxyprobe staining was concentrated in proximity to the vasculature, particularly areas adjacent to the central vein (Fig.3A.iv) as well as close to portal vein (Fig.3A.v). More intense staining was observed in areas in which the centrilobular mouse hepatocytes converge onto the central vein, as depicted by the darker brown patch next to the central vein, across the patchy staining pattern. This suggests that in the liver of Py17XL infected mice, greater hypoxic stress occurs adjacent to the central vein, particularly surrounding centrilobular mouse hepatocytes.

In the kidney, no hypoxyprobe staining was detected in any of the uninfected samples (Fig.3B.i, ii, iii). Hypoxyprobe staining in the Py17XL group was primarily seen in the glomeruli and the cortical tubules surrounding the glomeruli (Fig.3B.iv), as well as the tubules surrounding the renal vessel (Fig.3B.v), suggesting that within the kidney, the cortical tubules and glomeruli experience more hypoxic stress, during Py17XL infection.

No hypoxyprobe staining was observed in the small intestine of uninfected mice (Fig.3C.i, ii, iii), whereas Hypoxyprobe staining was apparent in the villi and the duodenum in Py17XL infected mice (Fig.3C.iv, v, vi). The lamina propria of the villi was found to have the most staining (Fig.3C.iv & vi). This suggests that within the small intestine, the lamina propria within the villi are the most susceptible to hypoxic stress, upon Py17XL infection.

### Tissue hypoxia in Py17XL leads to increased glycolysis

The transcription factor HIF-1α orchestrates the tissue response to hypoxia. To investigate whether tissue hypoxia in malaria triggers a HIF-1α-mediated response – specifically a glycolytic shift – the expression of *Hif-1α* and its downstream target genes involved in anaerobic glycolysis (glucose transporter 1, *Glut1*; lactate dehydrogenase A, *Ldha*; monocarboxylate transporter 4, *Mct4*) was analysed by RT-qPCR in the liver and kidney from uninfected control, PHZ-treated, and PY17XL-infected mice (Fig.4A, B). No significant changes in *Hif-1α* mRNA levels were detected in either the liver or kidney of the PHZ and Py17XL groups compared to uninfected control mice (Fig.4A, B), consistent with predominant post-translational regulation of HIF-1α activity. However, mRNA levels of downstream glycolytic target genes were significantly elevated in the liver of both the PHZ and Py17XL groups. *Glut1*, responsible for cellular glucose uptake, and *Mct4*, involved in lactate export, were more significantly upregulated in the Py17XL group compared to the PHZ group (Fig.4A). In the kidney, *Mct4* gene expression showed similar levels of upregulation in both the PHZ and Py17XL groups (Fig.4B), but *Glut1* and *Ldha* were only upregulated in PHZ treated mice (Fig.4B).

**Figure 4.**
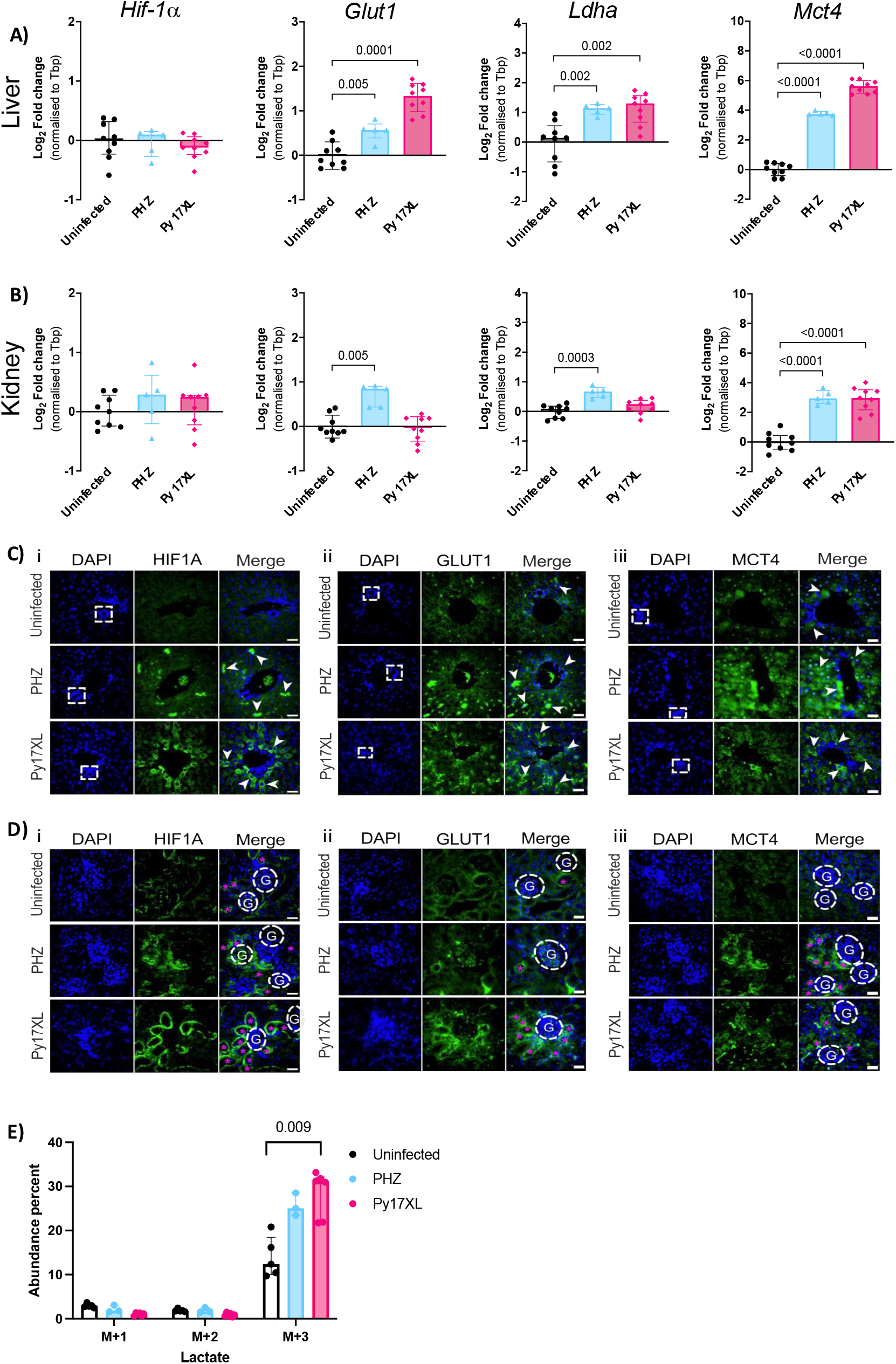
Tissue hypoxia leads to increased glycolysis in Py17XL infected mice. Expression of HIF-1α downstream target genes associated with anaerobic glycolysis measured by RT-qPCR in the (A) liver and (B) kidney from uninfected (n = 9), PHZ-treated (n = 5), and Py17XL infected mice (n = 9). Error bars show mean with standard deviation. Welch’s ANOVA test *p-value* <0.0001, *p-values* for Dunnett’s T3 multiple comparison test shown on the graph. *Hif-1α*, hypoxia-inducible factor 1 alpha subunit; *Glut1*, glucose transporter 1; *Ldha*, lactate dehydrogenase a; *Mct4*, monocarboxylate transporter 4. Immunofluorescence detection of HIF-1α expression in liver (C. i) and kidney (D. i); GLUT1 expression in liver (C. ii) and kidney (D. ii); and MCT4 expression in liver (C. iii) and kidney (D. iii) sections from uninfected (n = 3), PHZ-treated (n = 3), and Py17XL infected mice (n = 4). White dashed squares in the DAPI panel of the liver indicate the hepatic artery near a branch of the hepatic portal vein. Arrowheads in the liver sections point to cells or structures expressing the target protein. Each white circle in the kidney represents a glomerulus, while magenta asterisks indicate renal tubules with positive target protein staining. White arrows point to renal epithelial cells expressing GLUT1. Scale bar: 25 μm. RV, renal vessel; G, glomerulus. The percentage of the total abundance made up by each labeled isotopologue is shown for; E) Lactate, Kruskal-Wallis test *p-value* =0.001, *p-values* for Dunn’s multiple comparison test shown on the graph. Uninfected (n = 5), PHZ-treated (n = 3), Py17XL infected mice (n = 6). The different isotopologue species are indicated on the x-axes by M+(x), where x= the number of carbon atoms. In these graphs, the remaining percent is made up of the unlabeled M+0. Error bars show median with interquartile range.

These results demonstrate tissue-specific differences in the expression of *Hif-1α* induced glycolytic genes, with significant upregulation observed in the liver and kidney under hemolytic anemia and severe malaria infection.

To complement the analysis of changes in gene expression, the protein expression pattern of HIF-1α in liver and kidney of uninfected control, PHZ and Py17XL infected mice were examined through immunofluorescence staining (Fig.4C, D). Liver sections from the uninfected group did not exhibit any significant HIF-1α staining, while the PHZ and Py17XL groups showed enhanced expression of HIF-1α, indicating tissue hypoxia (Fig.4C.i). Different hepatic HIF-1α expression patterns were observed between the PHZ and Py17XL groups. In PHZ induced hemolytic anemia, HIF-1α was mainly expressed in hepatic sinusoids near the main hepatic vessels – the branch of the hepatic portal vein and the central vein (Fig.4C.i). This pattern of expression is similar to the areas showing intense hypoxyprobe staining, suggesting a correlation between tissue hypoxia and HIF-1α expression in these regions (Fig.3A). In the kidney, HIF-1α expression was primarily observed in renal tubules around the glomeruli across all groups with Py17XL-infected kidneys showing the highest expression (Fig.4D.i). Consistent with the HIF-1α patterns, immunofluorescence staining showed increased GLUT1 expression in the liver (Fig.4C.ii) and kidneys (Fig.4D.ii) of both PHZ and Py17XL groups compared to the uninfected group. In the kidney, GLUT1 expression was more distinct in renal tubules near glomeruli in Py17XL-infected mice compared to PHZ and uninfected groups (Fig.4D.ii). MCT4, another HIF-1α downstream target, exhibited distinct expression patterns in liver and kidney tissues. In the liver, PHZ-treated mice showed significantly stronger MCT4 expression in periportal hepatocytes, whereas Py17XL-infected mice showed MCT4 staining primarily in centrilobular hepatocytes (Fig.4C.iii). In the kidney, MCT4 was notable around glomeruli and renal vessels in PHZ-treated mice and Py17XL-infected mice (Fig.4D.iii).

Overall, these findings suggest that while both PHZ-induced hemolytic anemia and severe malaria trigger hypoxia and glycolytic responses, the affected tissues and regions within the liver and kidney differ between the two conditions, suggesting the distinct metabolic adaptations to hypoxia in each condition.

To assess if indeed more lactate is produced through glycolysis, we administered the stable isotope ^13^C_6_-labeled D-glucose and examined its systemic conversion to labeled lactate after 20 minutes (26) in plasma from Py17XL infected, PHZ treated and uninfected mice (Fig.4E). We found a significantly higher percent abundance of M+3 labeled lactate in Py17XL infected mice compared to uninfected mice, indicative of increased systemic glycolysis (Fig.4E).

### Liver and kidney dysfunction

Plasma alanine transaminase (ALT) and aspartate transaminase (AST) levels were measured to assess liver damage in malaria-infected (Py17XL and Py17XNL) and anemic (PHZ and anti-TER119) mice compared to uninfected controls. ALT levels were significantly elevated in both Py17XL (most notably) and Py17XNL-infected mice relative to uninfected controls (Fig.5A). For AST, a significant increase was observed only in Py17XL-infected mice (Fig.5B). While there was no significant elevation of liver enzymes in the non-infectious anemia models (Fig.5A, B). To investigate whether reductions in lactate and parasitemia following treatment correlated with liver enzyme recovery, we also measured AST and ALT levels after artesunate or combined artesunate and whole blood treatment in Py17XL infected mice. Both AST and ALT levels decreased post-treatment in these mice (Fig.5A and 5B).

**Figure 5.**
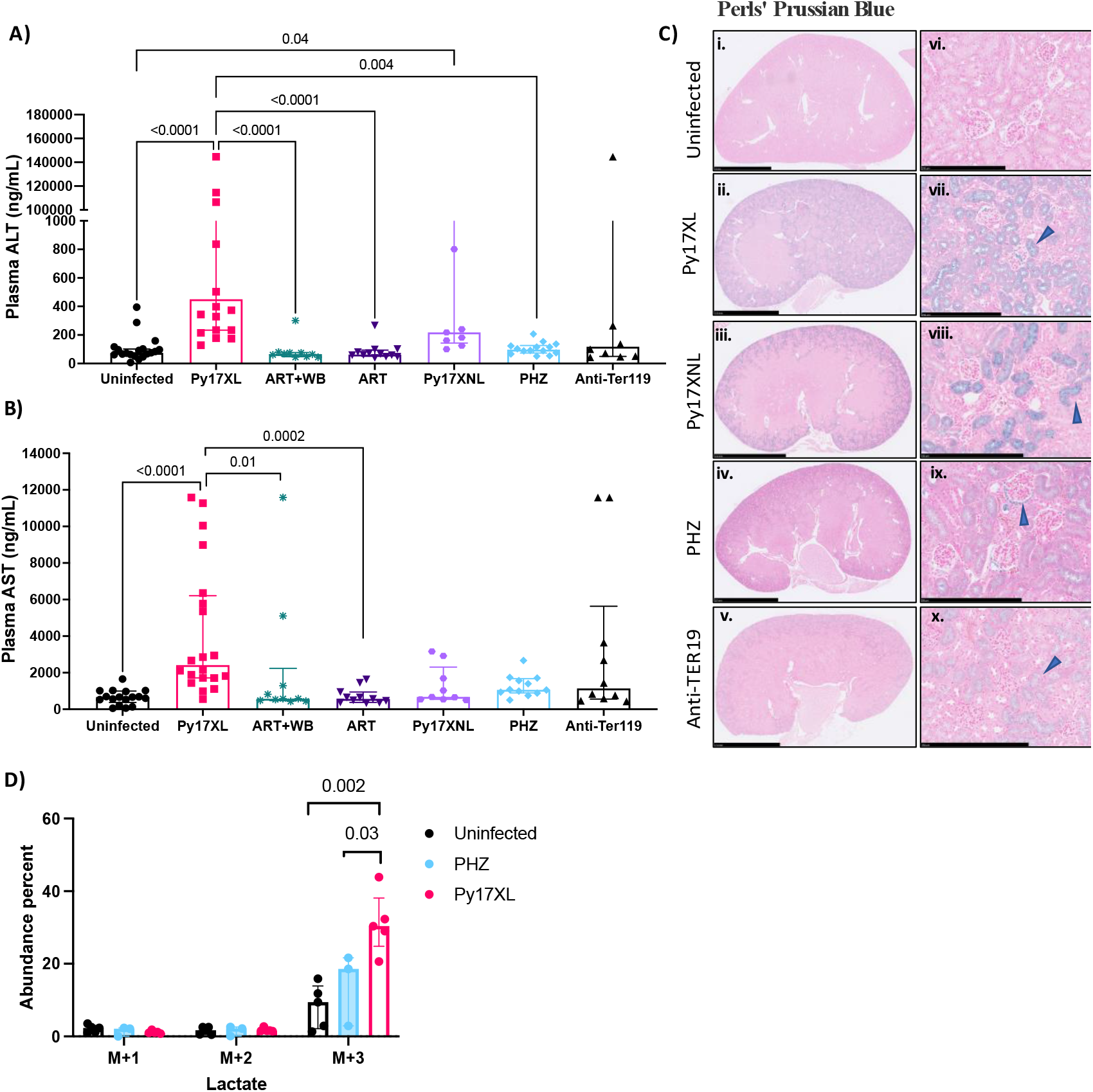
Liver and kidney dysfunction lead to reduced lactate clearance. A) Mouse plasma ALT levels (ng/ml) Kruskal-Wallis test *p-value* <0.0001, *p-values* for Dunn’s multiple comparison test shown on the graph, Uninfected n=19, Py17XL n=20, Py17XL+Art n=12, Py17XL +Art + WB n=10, Py17XNL n= 9, PHZ n=14, anti-TER119 n=10, Error bars show median with interquartile range. B) Mouse plasma AST levels (ng/ml), Kruskal-Wallis test *p-value* <0.0001, *p-values* for Dunn’s multiple comparison test shown on the graph, Uninfected n=17, Py17XL n=20, Py17XL+Art n=11, Py17XL +Art + WB n=10, Py17XNL n= 9, PHZ n=12, anti-TER119 n=10, Error bars show median with interquartile range. C) Perls’ Prussian Blue stain in mouse kidney. i-v: transverse kidney image, scale bar:2.5mm, vi-x: zoomed in images, scale bar:250um, blue arrowheads point towards hemosiderin deposits. Images shown representative of: Uninfected n=6, Py17XL n=8, Py17XNL n= 5, PHZ n=4, anti-TER119 n=4, D) The percentage of the total abundance made up by each labeled isotopologue is shown for lactate. One-way ANOVA *p-value=*0.003, *p-values* for Tukey’s multiple comparison test shown on the graph. Uninfected (n = 5), PHZ-treated (n = 3), Py17XL infected mice (n = 5). The different isotopologue species are indicated on the x-axes by M+(x), where x= the number of carbon atoms. In these graphs, the remaining percent is made up of the unlabeled M+0. Error bars show median with interquartile range. Eight to ten-week-old wild-type female C57BL/6J mice were used in all experiments.

Perls’ Prussian blue staining was used to assess renal iron deposition and inflammation associated with hemolysis. This revealed hemosiderin deposits in the renal proximal tubules and Bowman’s space in both malaria models (Py17XL and Py17XNL) (Fig.5C, ii, vii & iii, viii) and the anemic models (PHZ and anti-TER119 treated) (Fig.5C.iv, ix & v, x), compared to uninfected controls. Hemosiderin deposits were more pronounced in infected mice (Py17XL and Py17XNL).

To assess if lactate clearance is impaired due to liver and kidney damage, we administered the stable isotope ^13^C_3_-labeled sodium-L-Lactate and examined its systemic clearance after 20 minutes (27) in plasma from Py17XL infected, PHZ treated and uninfected mice (Fig.5D). We found a significantly higher percent abundance of M+3 labeled lactate in Py17XL infected mice compared to uninfected mice, indicative of decreased systemic lactate clearance (Fig. 5D).

These findings suggest that dysfunction in both liver and kidney of malaria infected mice can impair the systemic clearance of lactate.

### Trefoil factor 3 (TFF3) a biomarker of intestinal injury correlates with lactate and other severity malaria markers in human malaria

Trefoil factor 3 (TFF3), is a secreted protein in the small intestine and colon produced by goblet cells important for the maintenance of mucosal integrity and mucosal repair (28). Using the malaria SCRIPT (Search for Correlates of Recovery in the Patient Transcriptome) study subjects, we found that TFF3 is higher in severe malaria cases compared to uncomplicated malaria (Fig.6A). We also found a significant positive correlation between TFF3 and lactate (R = 0.32, *p-value* = 0.002) (Fig.6B) as well as TFF3 and *Plasmodium falciparum* histidine rich protein 2 (*Pf*HRP2) (R = 0.23, *p-value* = 0.032) (Fig.6C), and a significant negative correlation between TFF3 and hemoglobin (R = -0.21, *p-value* = 0.039) (Fig.6D). These findings suggest there is a link between gut dysfunction and severe malaria features such as hyperlactatemia, increased parasite load and anemia.

**Figure 6.**
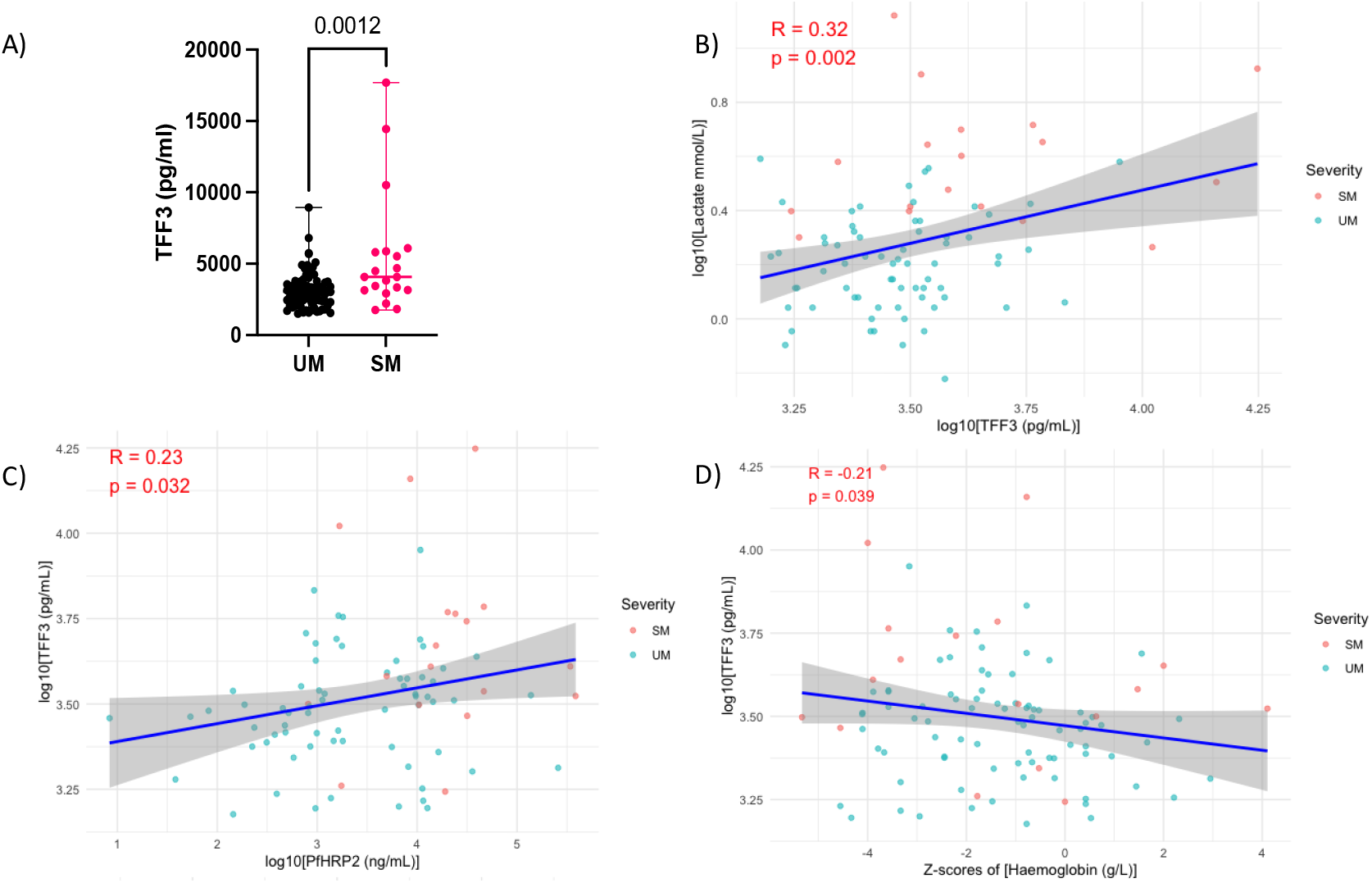
Trefoil factor 3 (TFF3) a biomarker of intestinal injury correlates with severity malaria markers in human malaria. A) Serum levels of TFF3 in severe (n=20) and uncomplicated (n=83) malaria patients. Mann-Whitney test *p-value*=0.0012. B) Pearson correlation between lactate and TFF3 (n=86). TFF3 expressed in pg/mL, and lactate expressed in mmol/L. The scatter plot shows the log10-transformed values of lactate (mmol/L) and TFF3 (pg/mL), with the blue line representing the linear regression fit (R = 0.32, *p-value* = 0.002). C) Pearson correlation between PfHRP2 and TFF3 (n=85). TFF3 levels expressed in pg/mL, and PfHRP2 levels expressed in ng/mL. The scatter plot shows the log10-transformed values of PfHRP2 (ng/mL) and TFF3 (pg/mL), with the blue line representing the linear regression fit (R = 0.23, *p-value* = 0.032). D) Pearson correlation between TFF3 and hemoglobin (n=100). Hemoglobin expressed in g/L, and TFF3 expressed in pg/mL. The scatter plot shows the log10-transformed values of TFF3 (pg/mL) and age-sex based z-scores of hemoglobin (g/L), with the blue line representing the linear regression fit (R = -0.21, *p-value* = 0.039). The data points are coloured based on disease severity, with red representing severe malaria (SM) and blue representing uncomplicated malaria (UM). The shaded area around the regression line indicates the 95% confidence interval.

## Discussion

This study provides important insights into the multifactorial drivers of hyperlactatemia in severe malaria, particularly emphasizing anemia as a key contributor while exploring other infection-specific mechanisms. Both the lethal (Py17XL) and non-lethal (Py17XNL) malaria mouse models exhibited significant hyperlactatemia at the nadir of hematocrit, supporting the notion that severe anemia is a central factor in elevated lactate levels. Notably, unlike *Plasmodium falciparum* infections in humans, where parasite sequestration in the microvasculature is thought to play a major role, our mouse models showed high lactate levels without the need for parasite sequestration. This suggests that parasite sequestration may not be an essential mechanism in the development of hyperlactatemia.

Our data further highlight the efficacy of combined artesunate and whole blood transfusion therapy, which significantly accelerated the resolution of both parasitemia and lactate levels compared to artesunate alone in Py17XL infected mice. This finding aligns with a recent study in experimental cerebral malaria (*P*.*berghei* ANKA infected mice) which showed that mice that received a combination of artesunate and whole blood as a single dose late in infection had a 90% survival rate compared to 54% in those who received artesunate alone due to recovery of platelets, hematocrit, angiopoietins and the integrity of blood brain barrier (29). Hematocrit in the combination treatment group was significantly higher compared to untreated Py17XL mice suggesting that whole blood transfusion corrected anemia. Understanding how whole blood synergistically works with artesunate to improve outcome and identifying which parameters recover would be crucial and informative for tailored human severe malaria treatments.

Our Hypoxyprobe staining results demonstrated distinct tissue hypoxia in the liver, kidneys, and small intestine, implicating these organs as key sites of oxygen deprivation during severe malaria. Using RT-qPCR we identified that tissue hypoxia leads to increased glycolysis in the liver and kidney shown by the increased expression of *Hif-1a* downstream target genes *Glut1, Ldha*, and *Mct4*. Tissue hypoxia, evidenced by increased protein expression of HIF-1α and GLUT1, LDHA and MCT4, was observed not only in malaria-infected mice but also in non-infectious anemia models, suggesting that anemia drives a metabolic shift towards glycolysis across conditions. The significantly elevated labeled plasma lactate in Py17XL-infected mice confirmed that increased glycolytic flux contributes substantially to hyperlactatemia under hypoxic conditions.

Previous studies have suggested that liver and kidney dysfunction found in severe malaria is responsible for reduced lactate clearance (6, 17), however there hasn’t been any experimental evidence to prove this. We found elevated levels of liver enzymes ALT and AST (significantly high only in Py17XL), hemozoin deposits, white blood cell infiltration and hemosiderin deposits (Perls’ Prussian Blue) in the kidneys of malaria infected mice, all indicative of tissue malfunction. In the anemia mouse models the levels of liver enzymes were not significantly increased compared to control uninfected mice however white blood cell infiltration in the liver was noted and anti-TER119 livers presented with areas of tissue necrosis. Perls’ Prussian Blue staining in the kidneys of anemic mice revealed hemosiderin deposits in the proximal renal tubules concordant with their hemolysis. While Py17XL-infected mice exhibited elevated ALT and AST levels, indicating liver damage, the non-infectious anemia models did not show significant increases in these enzymes. Iron and hemozoin deposits along with white blood cell infiltration in the liver and kidneys of infected mice suggest that tissue injury, likely driven by hemolysis and iron accumulation, may impair organ function. The hemosiderin accumulation, particularly in the kidneys, aligns with previous studies linking hemolytic stress to tissue fibrosis and senescence (30). The liver and kidney dysfunction also contributes to the reduced systemic clearance of ^13^C_3_-lactate in Py17XL infected mice compared to uninfected and PHZ treated mice.

Human studies in both adults and children have suggested that gut microvascular obstruction due to parasite sequestration in the gut microvasculature is responsible for gut dysfunction and permeability in severe malaria (31, 32). A recent study found that TFF3, a biomarker of intestinal injury, was increased in severe malaria pediatric cases compared to community control serum samples and correlated with mortality, acidosis, AKI and endothelial activation. Since we identified the presence of hypoxia in the gut (small intestine) of Py17XL infected mice we then wanted to examine gut dysfunction and its correlation to lactate and other severe malaria biomarkers using the SCRIPT study cohort including both severe and uncomplicated malaria patients. We found that TFF3 was significantly higher in severe malaria patients compared to uncomplicated malaria indicative of gut dysfunction associated with malaria severity. We also found significant associations with lactate, parasite burden, and anemia, suggesting that intestinal integrity is compromised during severe malaria. This provides further evidence that gut injury may be a contributing factor to the metabolic derangements observed in severe malaria, including hyperlactatemia.

In conclusion, this study provides clear evidence that anemia is a major driver of hyperlactatemia in malaria, but additional factors, including tissue hypoxia and organ dysfunction, contribute to the elevated lactate levels observed in severe cases. These findings suggest that therapies targeting both the correction of anemia and the mitigation of organ dysfunction, particularly in the liver, kidneys, and gut, may offer a more comprehensive approach to managing severe malaria. Future studies will aim to elucidate the specific mechanisms of lactate clearance and further refine treatment strategies for better clinical outcomes.

## Methods

### Animals and procedures

#### Ethics and approvals

Eight to ten-week-old, pathogen-free, female wild-type C57BL/6J mice were supplied by Charles River Laboratories. All mice were specified pathogen-free, housed in groups of five in individually ventilated cages, and provided ad libitum access to food and water. All experimental protocols and procedures were approved by the Imperial College Animal Welfare and Ethical Review Board, in accordance with the Laboratory Animal Science Association’s guidelines for good practice. Prior to any experimental interventions, all mice were acclimatized to the animal facility for one week.

#### Malaria mouse models

*Plasmodium yoelii* strains (*P. yoelii* 17XL [lethal] and *P. yoelii* 17XNL [non-lethal]) were serially passaged through C57BL/6J mice, following previously described protocols (23). Blood was collected via aseptic cardiac puncture from infected donor mice under non-recovery isoflurane anesthesia and diluted in sterile phosphate-buffered saline to achieve the desired parasite concentration. Experimental mice (eight to ten-week-old female C57BL/6J) were subsequently infected with 10^6^ live parasites via intraperitoneal injection.

Tail capillary blood samples were collected to prepare blood smears for parasitemia assessment and lactate measurement using the Lactate Pro 2 meter (HAB Direct). Parasitemia was quantified through microscopy of thin blood smears stained with 10% Giemsa, following established protocols (23). Heparinized blood was collected via cardiac puncture under non-recovery isoflurane anesthesia, and plasma was separated by centrifugation at 4,000g for 10 minutes. Hematocrit (%) were measured using capillary tubes (Ningbo Trustlab Instruments Co., Ltd, China) centrifuged at 900 x g for 10 minutes using a microhematocrit centrifuge (Servoprax, Germany). After centrifugation, the packed volume of the RBCs was measured.

#### Anemia mouse models

For the hemolytic anemia mouse model, eight to ten-week-old female C57BL/6J were treated with two doses, 24 hours apart, of 60 mg/kg phenylhydrazine (Sigma-Aldrich, UK) in a final volume of 200 μL through intraperitoneal injection (24). Blood lactate concentration was monitored daily upon the first dose of phenylhydrazine. At day 4 upon the first phenylhydrazine administration, all mice were euthanised via exsanguination by cardiac puncture under terminal isoflurane anaesthesia. For the extravascular hemolysis mouse model, eight to ten-week-old female C57BL/6J were treated with two doses of InVivoMAb anti-mouse TER119 (Bio X Cell) 1ug/g of body weight. First dose on day 0 and second dose on day 3 of experiment (25). As an isotype control for anti-TER119, 3 mice receive the same concentration of InVivoMAb rat IgG2b isotype control, anti-keyhole limpet hemocyanin (Bio X Cell). At day 4 upon the first anti-TER119 dose all mice were euthanised via exsanguination by cardiac puncture under terminal isoflurane anaesthesia. In both anemia mouse models tail capillary blood samples (1ul) were collected daily for lactate measurement using the Lactate Pro 2 meter (HAB Direct).

#### Organ collection and storage

Organs collected from *P. yoelii* 17XL, *P. yoelii* 17XNL, PHZ and anti-TER119 treated mice and fixed in 4% paraformaldehyde for 48 hr before being processed, snap frozen or fixed in RNAlater. Organs fixed in 4% paraformaldehyde were then paraffin embedded, cut, and stained with H&E and Perls’ Prussian Blue at IQPath (Institute of Neurology, University College, London, UK). Digitized images were taken (LEICA SCN400, Leica Microsystems UK) at IQPath (Institute of Neurology, University College, London, UK). Images were then viewed and examined with Aperio ImageScope software (Leica Biosystems Imaging, Inc).

#### Artesunate and combination of artesunate plus whole blood treatment in Py17XL mice

*P. yoelii* 17XL infected mice with similar levels of parasitemia were randomly allocated to artesunate or artesunate plus whole blood transfusion treatment groups on day 5 post-infection. Whole blood for transfusion was collected from healthy eight to ten-week-old female C57BL/6J mice via cardiac puncture under non-recovery isoflurane anesthesia. For artesunate treatment, mice received intraperitoneal injection of 30 mg/kg artesunate (A3731, Sigma-Aldrich) on day 5 and 6 post-infection. For the artesunate plus whole blood transfusion group, mice were treated with 30 mg/kg artesunate via intraperitoneal injection, followed by 250 μL whole blood transfusion through intraperitoneal injection on day 5 and 6 post-infection. On day 7 post-infection, all mice were euthanized via exsanguination by cardiac puncture under terminal isoflurane anaesthesia.

#### Administration of ^13^C_6_-labeled D-glucose or ^13^C_3_-labeled Sodium L-lactate in Py17XL, PHZ and uninfected mice

*P. yoelii* 17XL infected mice (at the humane endpoint), PHZ treated (at peak anemia) and control uninfected mice were injected intraperitoneally with either 1 g/kg of ^13^C_6_-labeled D-glucose (CLM-1396-1, Cambridge Isotope Laboratories) dissolved in sterile PBS or 0.5 mg/gram of body weight ^13^C_3_ Sodium L-lactate (CLM-1579-0.5, 20% w/w in water, Cambridge Isotope Laboratories). Twenty minutes after injection mice were euthanized via exsanguination by cardiac puncture under non-recovery isoflurane anesthesia. Plasma was collected for liquid chromatography-mass spectrometry.

#### Hypoxyprobe

Py17XL infected mice (at the humane endpoint) and control uninfected mice (eight to ten-week-old) were used for *in vivo* Hypoxyprobe administration. Each mouse was injected intraperitoneally with 2 mg of Hypoxyprobe (pimonidazole-HCl) (Hypoxyprobe Inc, USA) dissolved in 200 μL of sterile saline solution 1.5 hours before euthanasia. This timing was selected to ensure optimal tissue uptake of the Hypoxyprobe (33). Following the injection, all mice were euthanised by cardiac puncture under non-recovery isoflurane anaesthesia, and organs (liver, kidney, brain, spleen, gut, tail snip, heart and skeletal muscles) were collected immediately and placed in 4% paraformaldehyde solution at 4°C for 48 hours for fixation. After fixation, the organs were transferred to PBS at 4°C until further processing. This was followed by paraffin embedding and tissue sectioning performed by IQ Path (UCL Queen Square Institute of Neurology). Paraffin-embedded tissue sections from the liver, kidney, lungs, spleen, heart, tail, gut, and brain were deparaffinized in xylene, followed by rehydration in graded ethanol. Antigen retrieval was performed using Uni-TRIEVE Mild Temperature Universal Retrieval Solution (Innovex Biosciences, UK) at 70°C for 1 hour. Blocking was performed using BLOXALL® Endogenous Blocking Solution, Peroxidase and Alkaline Phosphatase (2BScientific, UK) for 10 minutes, followed by washing in PBS. Sections were then incubated with M.O.M.® (Mouse on Mouse) Immunodetection Kit diluent (Vector Laboratories, USA) at room temperature for 1 minute, followed by incubation with primary antibody anti-pimonidazole (Mab1) (Hypoxyprobe Inc, USA) at a 1:200 dilution in M.O.M. diluent, overnight at 4°C. M.O.M. biotinylated anti-mouse IgG was applied for a 15-minute incubation, followed by PBS washing. Next, sections were incubated with Vectastain ABC solution (Vector Laboratories, USA) followed by DAB substrate kit (2BScientific, UK). The sections were counterstained with haematoxylin and then dehydrated in ethanol and xylene before mounting with VectaMount solution (Vector Laboratories, USA) and coverslip. Digitized images were taken (LEICA SCN400, Leica Microsystems UK) at IQPath (Institute of Neurology, University College, London, UK). Images were then viewed and examined with Aperio ImageScope software (Leica Biosystems Imaging, Inc).

#### Immunofluorescence on mouse organs

Paraffin-embedded tissue sections from control, PHZ treated and Py17XL infected mice were deparaffinised by immersing in xylene and then hydrated with graded ethanol. Antigen retrieval was done in Uni-TRIEVE Mild Temperature Universal Retrieval Solution (Innovex biosciences, UK) at 65 °C for 45 min. To prevent any non-specific staining prior to antibody incubation, sections underwent blocking step with normal goat serum (Vector Laboratories, USA) at room temperature for an hour. After blocking, sections were incubated with listed primary antibodies at appropriate dilution in normal goat serum overnight at 4 °C (Table 1). Sections were then incubated with fluorochrome conjugated secondary antibody – goat anti-rabbit IgG H&L (Alexa Fluor® 488, ab150077, Abcam, UK) diluted in 1:500 ratio with PBS at room temperature for 30 min in the dark. After antibody incubation, slides were washed with PBST (0.1% Tween in 1X PBS) three times for 5 min each wash, followed by mounting with 4’,6-diamino-2-phenylindole (DAPI) containing medium which counterstains the nuclei (Vector Laboratories). Sections were then stored at 4 °C for imaging. Immunofluorescence imaging was conducted through an inverted Wide Field light-emitting diode (LED) illumination Axio observer 7 microscope (Zeiss, Germany) and the highly sensitive Hamamatsu Flash 4 camera at Imperial FILM facility. Through microscopy, 1,121 images were obtained from 12 mice. Images were then visualised through open source imaging software (Fiji).

**Table 1.**
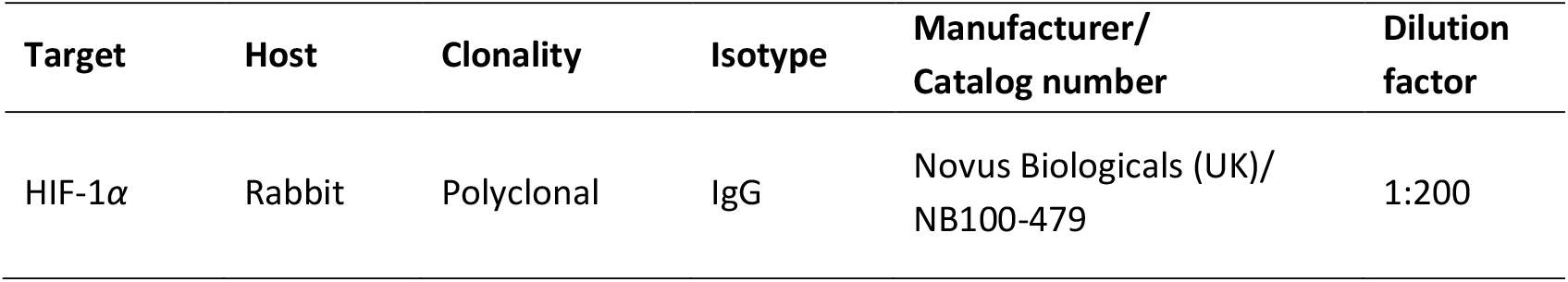
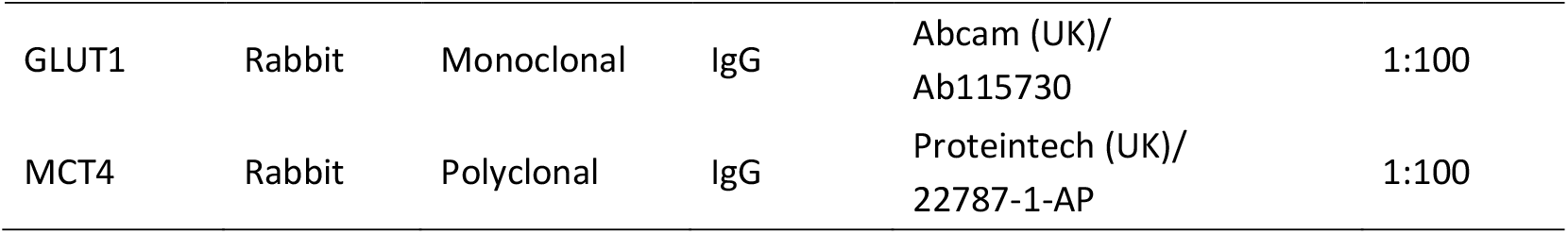
Primary antibodies used for immunofluorescence.

### Gene expression assay

#### Tissue homogenisation

Mouse liver, kidney, ear, and skeletal muscle stored at -80°C were thawed prior to tissue homogenisation. The amount of each tissue type for homogenisation was based on the manufacturer’s protocol. The tissue was placed in Lysing Matrix D 2 mL tubes (MP Biomedicals, USA), with 500 – 1000 μL buffer RLT (Qiagen, UK). Homogenisation was performed using the FastPrep-24TM 5G (MP Biomedicals) following the manufacturer’s recommended programs for each tissue type. The Lysing Matrix D tubes were centrifuged at 8000 x g at room temperature for 1 min.

#### RNA Extraction

200 – 500 μL of tissue lysates were transferred to new reaction tubes. An equal volume of 70% ethanol was then added and mixed. RNA extraction was performed using the RNeasy Mini Plus Kit (Qiagen) following the manufacturer’s instructions. RNA was eluted twice with RNase-free water (Thermo Fisher Scientific, UK). The total RNA yield was measured using a NanoDrop® ND-1000 Spectrophotometer (Thermo Fisher Scientific, UK).

#### Reverse Transcription

cDNA synthesis from the extracted RNA was processed by using the QuantiTect Reverse Transcription Kit (Qiagen, UK). Following the manufacturer’s recommended protocol, 1 μg of RNA was used for the reverse transcription. Each reaction tube contained an amount of RNA corresponding to 1 μg, adjusted with RNase-free water to a total volume of 14 μL. were placed in a Veriti™ Dx 96-well Thermal Cycler (Applied Biosystems, UK) and incubated at 42 °C for 2 minutes. Reverse transcriptase, random primers, and reverse transcription buffer were added to each tube. The thermal cycling conditions were set as follows: 42 °C for 20 minutes and 95 °C for 3 minutes. The cDNA samples were stored at -20 °C for subsequent qPCR experiments.

#### TaqMan RT-qPCR

TaqMan RT-qPCR was performed using 2x TaqMan® Fast Advanced Master Mix (Applied Biosystems, UK) and 20x mouse-specific TaqMan Gene Expression Assays (Applied Biosystems, UK). Each target gene was analyzed using the corresponding TaqMan Gene Expression Assay, as detailed in Table 2. Duplex RT-qPCR was performed using a FAM dye-labelled TaqMan Gene Expression Assay for the target gene and a VIC dye-labelled TaqMan Gene Expression Assay for Tbp, which served as the endogenous reference control. Master mix was prepared following the manufacturer’s recommendations (Table 3). A total of 10 μL of reaction mixture was loaded into each well of a MicroAmpTM Fast Optical 96-well qPCR reaction plate (Applied Biosystems, UK) and centrifuged at 1500 rpm for 1 minute. The RT-qPCR reactions were carried out using a StepOnePlusTM Real-Time PCR System (Applied Biosystems, UK) following the manufacturer’s protocol (Table 4). Data analysis was performed using ThermoFisher Scientific StepOnePlusTM Software version 2.3. Relative gene expression was evaluated using the comparative cycle threshold (CT) method, assuming (i) 100% PCR efficiency and (ii) similar PCR efficiency between the target gene and the endogenous reference gene.

**Table 2.** TaqMan Gene Expression Assays.

| Gene | Assay ID | Probe type |
| --- | --- | --- |
| *Hif-1α* | Mm00468869_m1 | FAM-MGB |
| *Glut1 (Slc2a1)* | Mm00441480_m1 | FAM-MGB |
| *Ldha* | Mm01612132_m1 | FAM-MGB |
| *Mct4 (Slc16a3)* | Mm00446102_m1 | FAM-MGB |
| *Tbp* | Mm01277042_m1 | VIC-MGB |

**Table 3.** Master Mix composition.

| Component | Volume per reaction (μL) |
| --- | --- |
| TaqMan<sup>®</sup> Fast Advanced Master Mix (2X) | 5.0 |
| TaqMan<sup>®</sup> Gene Expression Assay (20X)<br>- Target gene (FAM-MGB) | 0.5 |
| TaqMan<sup>®</sup> Gene Expression Assay (20X)<br>- Endogenous reference gene *Tbp* (VIC-MGB) | 0.5 |
| Nuclease-Free Water | 3.0 |
| Total volume of Master Mix | 9.0 |
| cDNA template of each sample | 1.0 |
| Total volume per reaction | 10.0 |

**Table 4.** Thermal cycling settings on the StepOnePlus™ Real-Time PCR System.

| Steps | Cycles | Temperature | Time |
| --- | --- | --- | --- |
| UNG incubation | 1 | 50 °C | 2 min |
| Polymerase activation | 1 | 95 °C | 2 min |
| Denaturation | 40 | 95 °C | 1 sec |
| Annealing/ extension |  | 60 °C | 20 sec |

**Table 5.**
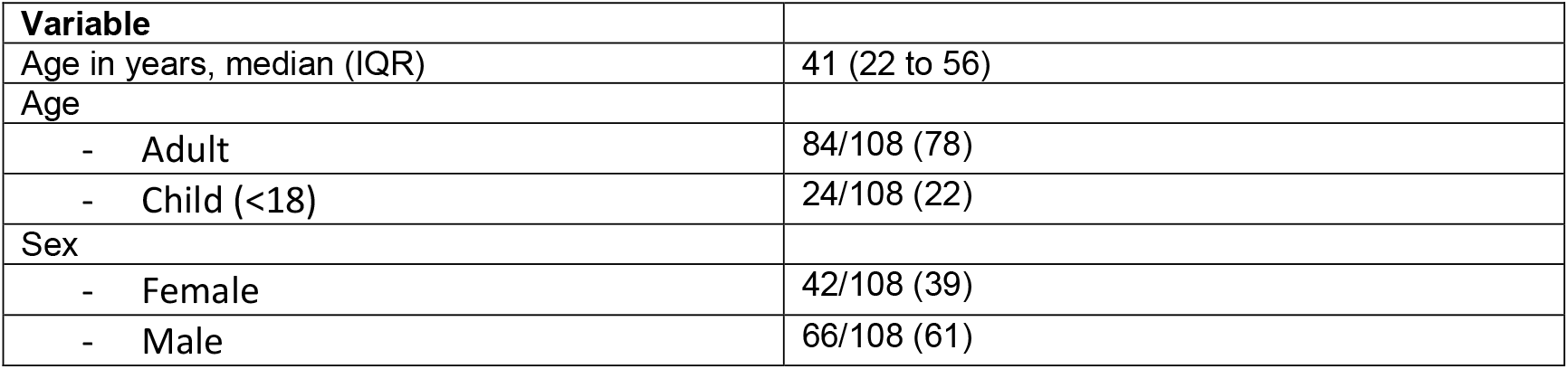
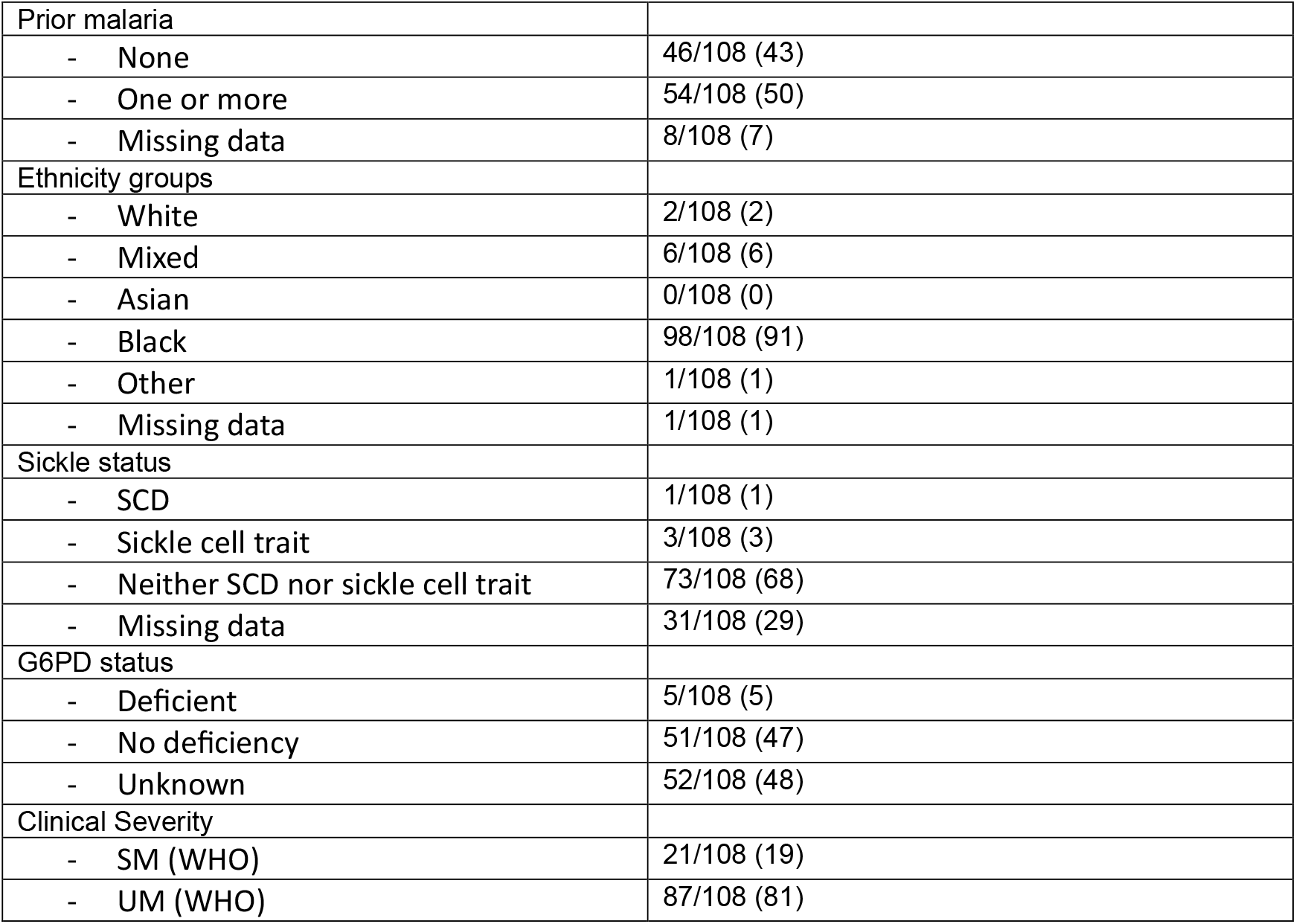
Demographic and clinical features of SCRIPT subjects used for the intestinal injury biomarker trefoil factor 3 Luminex assay.

| Variable |  |
| --- | --- |
| Age in years, median (IQR) | 41 (22 to 56) |
| Age |  |
| - Adult | 84/108 (78) |
| - Child (<18) | 24/108 (22) |
| Sex |  |
| - Female | 42/108 (39) |
| - Male | 66/108 (61) |
| Prior malaria |  |
| - None | 46/108 (43) |
| - One or more | 54/108 (50) |
| - Missing data | 8/108 (7) |
| Ethnicity groups |  |
| - White | 2/108 (2) |
| - Mixed | 6/108 (6) |
| - Asian | 0/108 (0) |
| - Black | 98/108 (91) |
| - Other | 1/108 (1) |
| - Missing data | 1/108 (1) |
| Sickle status |  |
| - SCD | 1/108 (1) |
| - Sickle cell trait | 3/108 (3) |
| - Neither SCD nor sickle cell trait | 73/108 (68) |
| - Missing data | 31/108 (29) |
| G6PD status |  |
| - Deficient | 5/108 (5) |
| - No deficiency | 51/108 (47) |
| - Unknown | 52/108 (48) |
| Clinical Severity |  |
| - SM (WHO) | 21/108 (19) |
| - UM (WHO) | 87/108 (81) |

CT values for target genes were normalised against the endogenous reference control Tbp. Gene expression analysis for all samples was based on technical duplicates.

#### ELISAs (ALT & AST)

Plasma levels of ALT and AST were measured following the manufacturer’s protocol using the Mouse ELISA Kit for ALT (ab282882, Abcam) and AST (ab263882, Abcam). The optical density was recorded using a microplate reader set at a wavelength of 450 nm.

#### Sample preparation for Liquid chromatography-mass spectrometry

During sample preparation for metabolomics, only LC-MS grade solvents were used and samples were kept on ice or stored at -70°C as required. Briefly, 40 μl from each plasma sample was added to 160 μl of ice-cold acetonitrile:methanol (1:1) and vortexed before addition of 200 μl of ice-cold acetonitrile:methanol:water (2:2:1). Samples were then vortexed again and then centrifuged at 15,000 x g for 10 minutes. Supernatant was taken and filtered using pre-washed SpinX 0.22 μm centrifuge filters (Corning 8161) by centrifuging at 15,000 x g for 10 minutes. For quality control purposes, a small aliquot of each sample filtrate was combined to create a pooled sample. This pooled sample was then used to create a dilution series which was run on the LC-MS with the samples; analysis of this dilution series was used to confirm that measured ion peaks were within linear range. Samples were diluted 1 in 4 in acetonitrile:methanol:water (2:2:1) prior to LC-MS analysis.

#### Liquid chromatography-mass spectrometry for stable isotope labeling analysis

The data were acquired with an Agilent 1290 Infinity II UHPLC coupled to a 6545 LC/Q-TOF system. Chromatographic separation was performed with an Agilent InfinityLab Poroshell 120 HILIC-Z (2.1 × 100 mm, 2.7 μm (p/n 675775-924)) column. Mobile phase A consisted of 10 mM ammonium acetate in water (pH 9) and mobile phase B consisted of 10 mM ammonium acetate (pH 9) in 15:85 (v:v) water/acetonitrile. Both mobile phases contained 5 μM Agilent InfinityLab Deactivator Additive (p/n 5191-4506). The following gradient was applied at a flow rate of 0.25 ml/min and a column temperature of 50°C: 0-2 minutes, 96% B; 5.5-8.5 minutes, 88% B; 9-14 minutes, 86% B; 17 minutes, 82% B; 23-24 minutes, 65% B; finishing with 5-minutes of re-equilibration at 96% B. Accurate mass spectrometry was performed using an Agilent Accurate Mass 6545 QTOF apparatus with the ESI ionization source operated in negative-ion mode. The following parameters were used: sheath gas temperature, 350°C; nebulizer pressure, 35 psig; sheath gas flow, 12 l min^-1^; capillary voltage, 3500 V; nozzle voltage, 0 V; and fragmentor voltage, 125 V. The data were collected in 4 GHz (high resolution) mode.

#### Data analysis for Liquid chromatography-mass spectrometry

Alignment of chromatograms, targeted feature extraction, and isotopologue analysis were performed using Agilent Profinder (version B.8.0.00, service pack 3). Metabolite identification was confirmed by comparison to standards.

### Human subjects

#### SCRIPT study

The SCRIPT (Search for Correlates of Recovery in the Patient Transcriptome) (Malaria) Study is a four-year observational cohort study aimed at identifying genetic and transcriptomic factors linked to recovery from malaria. It recruits patients of all ages with symptomatic *Plasmodium falciparum* malaria from selected hospitals in London. The primary goal is to identify genes whose expression correlates with a composite recovery score. Secondary goals include identifying genes tied to recovery rates for specific tissue or organ dysfunctions and understanding recovery pathways. Blood and urine samples are collected at multiple points post-diagnosis to track recovery markers.

#### Study Population

Patients were recruited from selected hospitals in London with a confirmed diagnosis of asexual stage parasitemia for any *Plasmodium* species. Patients with asymptomatic malaria, congenital malaria, *Plasmodium* gametocytemia without asexual parasitemia, or known HIV infection were excluded, as were those who had received antimalarial treatment in the 28 days prior to hospital presentation. For the analysis of organ damage biomarkers, only patients with *Plasmodium falciparum* infections were included, along with those whose antimalarial treatment began within 24 hours before or after the first blood sample collection. Patients co-infected with hepatitis B or C were excluded.

#### Ethics and Consent

The study received ethical approval from the Newcastle & North Tyneside 1 Research Ethics Committee (REC reference: 22/NE/0005). Informed consent was obtained from all participants or their legal representatives, with information sheets tailored to age groups. In cases where participants were unable to read, an impartial witness was present. Written consent was documented, and participants could withdraw from the study at any time, with the option to allow or revoke the use of previously collected data and samples. Where immediate consent was not feasible, a deferred consent approach was used, allowing for the collection of the first research sample alongside clinically indicated blood draws, with written consent sought thereafter.

#### Study Procedures

Upon blood film confirmation of asexual parasitemia, patients were approached for consent and enrolled in the study. Blood samples were collected at four different time points following diagnosis: timepoint 1 samples used here were collected as soon as possible after diagnosis. Sample collection aimed to capture dynamic trends in selected biomarkers indicative of tissue, organ, or organ system dysfunction.

#### Sample Collection, Processing, and Storage

BD Serum separating tubes (SST) were used to collect serum for biomarker analysis. Samples were stored at 4°C locally and transported to the Imperial College laboratory for processing. Blood samples in SST tubes were centrifuged at 2,000 x g for 10 minutes at 4°C to separate the serum. The serum was aliquoted and stored at -80°C until analysis.

#### TFF3 Intestinal Injury Biomarker Assay

The intestinal injury biomarker trefoil factor 3 (TFF3) was assessed in all frozen serum samples that met inclusion criteria. Luminex Human Discovery Assay (LXSAHM) (R&D Systems, Minneapolis, MN, United States) was used for the evaluation of TFF3 levels in serum samples from timepoint 1. Serum samples were centrifuged at 16,000 x g for 4 minutes before dilution. The samples were diluted as 1:6 using the diluent provided in the assay kit.

## Statistical tests

GraphPad Prism 10 (GraphPad Software) was used for statistical analyses of lactate concentration, parasitemia, HCT and ELISAs. The Shapiro-Wilk used to assess the normality of the data. Welch’s t-test, Welch’s ANOVA or ordinary one-way ANOVA was used for normally distributed data, while the Kruskal-Wallis test was applied to non-normally distributed data. Dunnett’s, Tukey’s or Dunn’s test was employed for post hoc multiple comparisons depending on the data distribution. For correlation analysis, Pearson’s correlation coefficient was used for normally distributed data. All tests were two-sided using a significance threshold of 5%. All experiments have been repeated at least twice to ensure reproducibility.

## Acknowledgments

We would like to thank the staff of Imperial College Central Biomedical Services for support with all animal experiments, the Facility for Imaging by Light Microscopy (FILM) at Imperial College London, and IQpath (Institute of Neurology, University College, London, UK) for their histology services. Infrastructure support for the SCRIPT study was provided by the NIHR Imperial Biomedical Research Centre (BRC).

